# Biogeographic and genomic signatures of thermal adaptation in facultative rhizobia

**DOI:** 10.1101/2025.01.14.632453

**Authors:** Chang-Yu Chang, Terrence Topping-Brown, Jazmine L. Rud, McCall B. Calvert, Gerardo Bencosme, Corlett W. Wood

## Abstract

Many plant endosymbionts are facultative, switching between host-associated and free-living stages. Extensive genomic and experimental studies suggest that adaptation during the saprophytic, off-host phase, rather than adaptation to hosts, primarily constrains the biogeographic distribution of these microbes. To test this hypothesis, we analyzed the growth capacities and genomic features of 38 *Sinorhizobium* strains nodulating *Medicago lupulina* (black medic), collected from two regions with distinct thermal environments. The warmer region is predominantly inhabited by *S. meliloti*, while *S. medicae* and non-symbiotic strains (lacking symbiosis genes, such as *S. canadensis* and *S. adherens*) are more common in the cooler region. Laboratory assays demonstrated that at 40°C, the upper temperature limit of their region of origin, *S. meliloti* remained viable, albeit with reduced growth, whereas *S. medicae* and non-symbiotic strains failed to grow under heat stress. Comparative genomics revealed isolation-by-distance in both the core and accessory genomes, particularly in *S. meliloti* in the warmer region, which exhibits less within-region thermal variation. This is consistent with an isolation-by-distance model where population divergence is governed by restricted gene flow. These findings suggest that metabolic constraints shape the regional distribution of this facultative microbial symbiont, while limited gene flow influences local population structure.

## INTRODUCTION

Understanding how microorganisms adapt to their environments across various life stages is fundamental to comprehending their ecological roles and evolutionary trajectories [1]. Many microbes are facultative symbionts, meaning that they switch between host-associated and free-living phases [2–4]. This dual lifestyle presents unique evolutionary challenges, as microbes must optimize their physiology and genetic makeup to succeed both within hosts and independently in the environment [5, 6]. The ability to navigate these contrasting niches influences not only their immediate survival but also their long-term geographic distribution and genetic diversity.

A key question in microbial ecology is what drives the biogeographic patterns observed in these facultative symbionts [7, 8]. While symbiont co-evolution with host plants or animals undoubtedly plays a role [9, 10], extensive experimental and genomic studies suggest that adaptation to abiotic environmental factors during free-living phases strongly influences their geographic distribution [9, 11–14]. Temperature, resource availability, and other environmental variables impose selective pressures that shape the genetic and phenotypic traits of these microbes, ultimately determining where they can thrive and persist [12, 13].

Two hypotheses predict how evolutionary and ecological processes operate across different spatial scales. At a regional level, metabolic constraints—such as thermal tolerance and resource utilization—are the primary factors limiting microbial distribution. This is supported by the abundant literature on the global biogeography of microbial community diversity [15, 16]. Under this hypothesis, individuals with enhanced thermal performance are more likely to establish in warmer regions, and we expect to observe differentiation in the genomic signatures of thermal stress response in populations distributed across different climates. In contrast, at finer spatial scales where abiotic gradients are less pronounced and selective pressures are more variable, gene flow becomes the main factor shaping local population structure [17]. Depending on the frequency of gene flow, local populations may either exhibit significant genetic differentiation or remain genetically similar. Such divergence can manifest not only in allelic variations of core genes but also in differences in accessory genome content [18].

Here we investigated the contribution of thermal tolerance to regional biogeographic patterns in nitrogen-fixing symbiotic bacteria known as rhizobia. Rhizobia are a prominent example of facultative symbionts that transition between a plant endosymbiotic phase and a free-living soil-dwelling phase [19, 20]. Rhizobia induce nodule formation in legume plants, and perform biological nitrogen fixation for their host [21]. Despite their ecological and agricultural importance as plant symbionts, much less is known about rhizobia’s saprophytic soil phase, during which their fitness depends on coping with environmental stresses and navigating microbial competition [19, 22, 23]. Like other soil microbes, rhizobia growth and survival during this free-living phase are sensitive to abiotic factors such as soil type and temperature [24–26]. Previous rhizobia biogeography has been shown to correlate with environmental conditions such as soil properties. For example, among soybean-nodulating rhizobia, *Sinorhizobium* predominates in alkaline soils, whereas *Bradyrhizobium* is more common in neutral or acidic soils [13, 14]. Understanding how free-living rhizobia adapted to abiotic environments is essential for explaining rhizobial biogeography, but assessments of growth performance across strains remain limited to representative species and strains [27].

Because of the challenges across the two life histories, rhizobia generally possess large genomes with key functions related to symbiosis and metabolism organized into genetic modules [28, 29]. Population genomics of *Sinorhizobium meliloti* across a large geographic range suggested a relatively more frequent gene flow in genomic regions relevant to symbiosis but not the conserved functions [29]. In addition, rhizobia possess a flexible pangenome with variation in the accessory gene content like many other prokaryotes [30]. This genomic architecture may result in population differentiation that reflects geographic and ecological influences that may manifest not only in the conserved core genes but also flexible accessory genomes [14]. However, variation in gene content along geographic distance is less explored in rhizobia.

In this study, we investigated the genomic feature and growth capacity in a set of *Sinorhizobium* strains that nodulate a widespread legume species, *Medicago lupulina*. We conducted laboratory assays to measure rhizobia growth at different temperatures and performed comparative genomic analyses of 38 strains’ core and accessory genomes. We found that these strains exhibit distinct biogeographic patterns across two regions with contrasting thermal features (Fig. S1): *S. meliloti* predominated in the warmer region, whereas *S. medicae* and non-symbiotic strains were more common in the cooler one (Fig. 1). Consistent with this observation, *S. meliloti* exhibited higher thermal tolerance in laboratory growth assays than *S. medicae*. Our results suggest that adaptation to thermal conditions in the free-living phase underpins the regional distribution of genus. By contrast, there was little evidence for thermal local adaptation with regions. Instead, within-region population differentiation is consistent with a pattern of isolation-by-distance in both the core and accessory genomes within species. These findings illuminate how ecological selection and dispersal processes intertwine to influence the biogeography of facultative symbionts.

**Figure 1.**
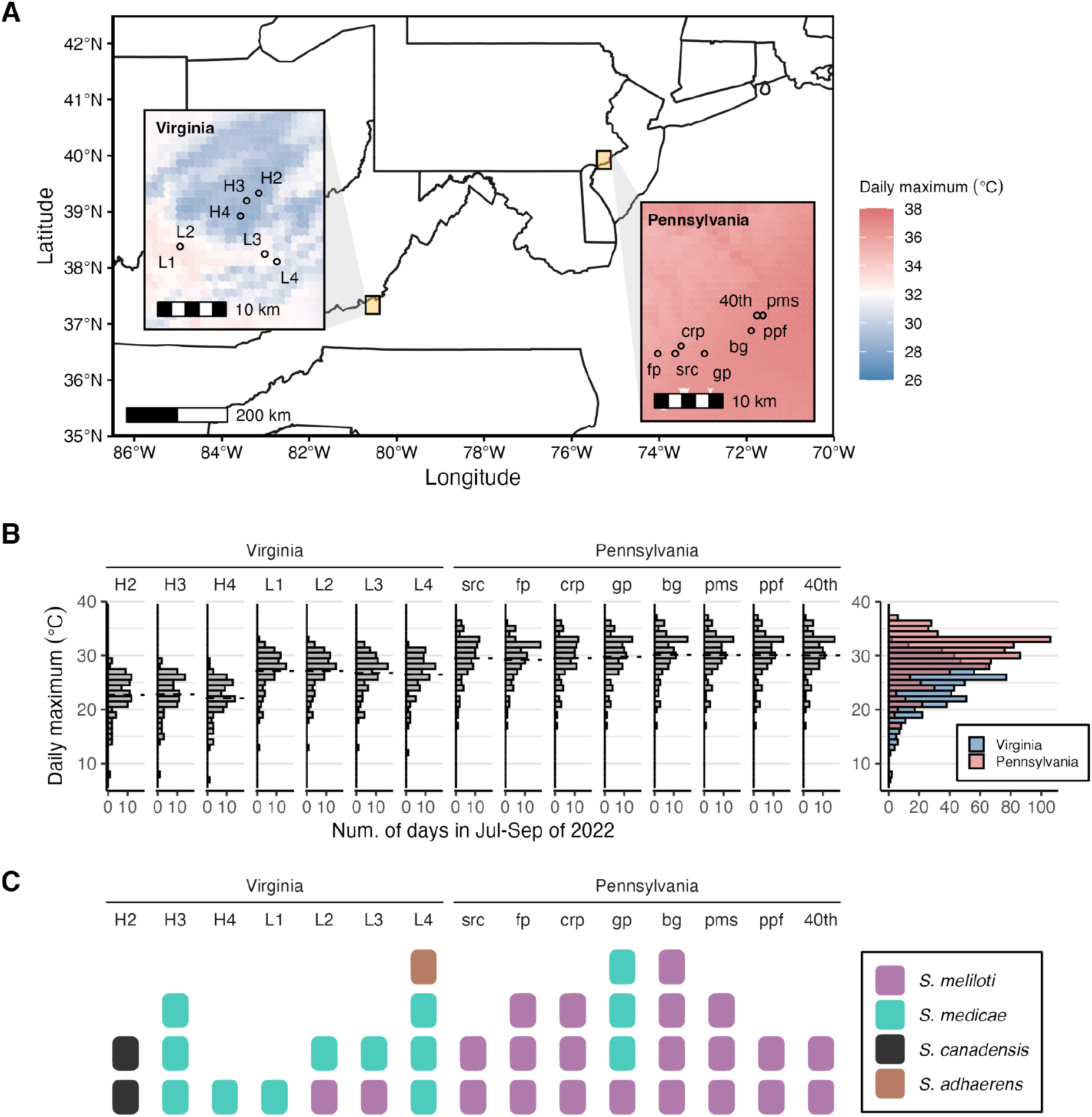
Sampling sites, thermal environments, and Sinorhizobium isolates. We sampled 38 *Sinorhizobium* strains from the nodules of *Medicago lupulina* that sampled from two geographic regions—Pennsylvania and Virginia—that differed in thermal environments. (A) The color represents the hottest daily maximum surface air temperature from July to September in the year of 2022 during which the samples were collected. Climate data was inter- and extrapolated from daily meteorological observations using Daymet [31] (SI section 1). Between Pennsylvania and Virginia, sampling sites are more than 500 km apart, whereas within regions the sites are 0.8-18 km apart. (B) Daily maximum of the sampling sites from July to September in the year of 2022. The dashed line represents the average at each site. The rightmost panel shows the accumulated counts across all sites. (C) The strains and species isolated from each sampling site.

## MATERIALS AND METHODS

### Study system

The *Sinorhizobium-Medicago* system is a model for investigating the evolution of symbiotic mutualism [32, 33]. *M. lupulina*, an annual or short-lived perennial weed native to Eurasia, was introduced to the Americas in the eighteenth century [34]. This species is a pioneer plant found in natural environments such as open grasslands, as well as in disturbed areas like lawns and urban green spaces [34]. In North America, *M. lupulina* is commonly associated with two symbiotic *Sinorhizobium* species: *S. meliloti* and *S. medicae* [35]. These two symbiotic species have a stable multipartite genome, comprising a chromosome (3.4-4.2 Mbp), a symbiosis megaplasmid pSymA (0.9-1.7 Mbp) that houses symbiosis genes essential for nodulation and nitrogen fixation (*nod, nif*, and *fix*), and a saprophytic chromid pSymB (1.6-2.0 Mbp) that contains chromosome-like and catabolic genes essential for saprophytic competence [28, 29, 33]. Many strains also possess smaller accessory plasmids (hereafter, pAcce) that encode a variety of functions [29]. The nonsymbiotic clade of *Sinorhizobium* is also composed of saprotrophic species such as the *S. adhaerens*, and *S. canadensis*, both of which have been reported to be isolated from root nodules despite lacking symbiosis plasmids [27, 36].

### Field sites and sampling

We sampled the root nodules of *M. lupulina* from 15 sites between July-September of 2022 from two geographic regions, including seven sites in the Appalachian Mountains in Giles County, Virginia (VA) and eight sites in the greater Philadelphia, Pennsylvania region (PA). Upon collection, we stored *M. lupulina* plants in zip-top bags at 4℃ for up to one week before we isolated the rhizobia strains from the nodules. We then isolated rhizobia with the following steps. First, we surface-sterilized nodules by submerging them in 100% ethanol for 20 seconds, followed by household bleach (5.25% sodium hypochlorite) for 20 seconds. We rinsed the nodules in distilled water and crushed them in a 20 μL droplet of phosphate buffer saline (PBS) on a Petri dish lid with flame-sterilized sharp tweezers. We then used a sterilized inoculation loop and streaked the inoculum on a solid tryptone-yeast (TY) agar plate, incubated the plates at 30℃ for three days until the visible colonies formed, and re-streaked morphologically distinguishable colonies on fresh TY agar plates. We repeated this re-streaking step three times until colonies in one agar plate had consistent morphologies. On the last day of isolation, we suspended the rhizobia colonies in 500 μL of PBS, mixed the suspension with an equal amount of 80% glycerol, and stored the mixture at -80℃ as stocks. The glycerol stocks were later used for whole-genome long-read sequencing, growth curve assays, and plant experiments. We obtained 38 distinct *Sinorhizobium* strains, including 15 strains from 7 sites in Virginia and 23 from 8 sites in Pennsylvania (Table 1).

**Table 1.** *Sinorhizobium* strains in this study. The “+” sign indicates whether a strain was used in the experiment: 14 inoculated to the host *M. lupulina* and 32 in the growth curve assay.

|  | Region | BLAST rRNA gene |  | BLAST genome |  | Strain |  | Experiment |  |
| --- | --- | --- | --- | --- | --- | --- | --- | --- | --- |
|  |  | Species | Identity (%) | Species | Identity (%) | Internal ID | Genome ID | Plant inoculation | Growth |
| 1 | Virginia | *S. fredii* | 98.9 | *S. adhaerens* | 99.0 | L4M2R2 | g15 | + | + |
| 2 |  | *S. fredii* | 98.8 | *S. canadensis* | 92.7 | H2M3R1 | g2 | + | + |
| 3 |  | *S. fredii* | 98.7 | *S. canadensis* | 91.4 | H2M3R2 | g3 |  | + |
| 4 |  | *S. meliloti* | 99.5 | *S. medicae* | 99.9 | H3M1R1 | g4 | + | + |
| 5 |  | *S. meliloti* | 99.5 | *S. medicae* | 99.9 | H3M3R2 | g5 |  | + |
| 6 |  | *S. meliloti* | 99.5 | *S. medicae* | 99.9 | H3M4R1 | g6 |  | + |
| 7 |  | *S. meliloti* | 99.5 | *S. medicae* | 99.9 | H4M5R1 | g8 | + | + |
| 8 |  | *S. meliloti* | 99.5 | *S. medicae* | 99.9 | L1M2R2 | g9 |  | + |
| 9 |  | *S. meliloti* | 99.5 | *S. medicae* | 99.9 | L2M4R1 | g11 |  | + |
| 10 |  | *S. meliloti* | 99.5 | *S. medicae* | 100.0 | L3M5R1 | g13 | + | + |
| 11 |  | *S. meliloti* | 99.5 | *S. medicae* | 99.8 | L4M3R3 | g16 |  | + |
| 12 |  | *S. meliloti* | 99.5 | *S. medicae* | 99.8 | L4M4R1 | g17 |  | + |
| 13 |  | *S. meliloti* | 99.5 | *S. medicae* | 100.0 | L4M7R1 | g19 |  | + |
| 14 |  | *S. meliloti* | 99.7 | *S. meliloti* | 99.6 | L2M2R1 | g10 | + | + |
| 15 |  | *S. meliloti* | 99.7 | *S. meliloti* | 99.9 | L3M1R1 | g20 |  | + |
| 16 | Pennsylvania | *S. meliloti* | 99.5 | *S. medicae* | 99.9 | gp1-2 | g29 |  | + |
| 17 |  | *S. meliloti* | 99.5 | *S. medicae* | 99.9 | gp1-3 | g30 |  | + |
| 18 |  | *S. meliloti* | 99.5 | *S. medicae* | 99.6 | gp1-1 | g40 | + | + |
| 19 |  | *S. meliloti* | 99.7 | *S. meliloti* | 99.9 | src-2 | g21 | + | + |
| 20 |  | *S. meliloti* | 99.7 | *S. meliloti* | 99.8 | fp1-2 | g22 | + | + |
| 21 |  | *S. meliloti* | 99.7 | *S. meliloti* | 99.8 | fp1-3 | g23 |  | + |
| 22 |  | *S. meliloti* | 99.7 | *S. meliloti* | 99.9 | fp2-2 | g24 |  | + |
| 23 |  | *S. meliloti* | 99.7 | *S. meliloti* | 100.0 | crp1-2 | g25 | + | + |
| 24 |  | *S. meliloti* | 99.7 | *S. meliloti* | 100.0 | crp1-3 | g26 |  | + |
| 25 |  | *S. meliloti* | 99.7 | *S. meliloti* | 100.0 | crp2-2 | g27 |  | + |
| 26 |  | *S. meliloti* | 99.7 | *S. meliloti* | 99.9 | src-2b | g39 |  |  |
| 27 |  | *S. meliloti* | 99.7 | *S. meliloti* | 99.9 | src-1 | g44 |  |  |
| 28 |  | *S. meliloti* | 99.7 | *S. meliloti* | 99.9 | bg-2 | g31 | + | + |
| 29 |  | *S. meliloti* | 99.7 | *S. meliloti* | 100.0 | bg-3 | g32 |  | + |
| 30 |  | *S. meliloti* | 99.7 | *S. meliloti* | 100.0 | pms-1 | g33 |  | + |
| 31 |  | *S. meliloti* | 99.7 | *S. meliloti* | 100.0 | pms-2 | g34 | + | + |
| 32 |  | *S. meliloti* | 99.7 | *S. meliloti* | 99.9 | pms-3 | g35 |  | + |
| 33 |  | *S. meliloti* | 99.7 | *S. meliloti* | 100.0 | ppf-1 | g36 |  | + |
| 34 |  | *S. meliloti* | 99.7 | *S. meliloti* | 100.0 | 40th-1 | g37 |  | + |
| 35 |  | *S. meliloti* | 99.7 | *S. meliloti* | 100.0 | bg-2b | g41 |  |  |
| 36 |  | *S. meliloti* | 99.7 | *S. meliloti* | 100.0 | ppf-2 | g42 | + |  |
| 37 |  | *S. meliloti* | 99.7 | *S. meliloti* | 100.0 | 40th-2 | g43 | + |  |
| 38 |  | *S. meliloti* | 99.7 | *S. meliloti* | 100.0 | bg-1 | g45 |  |  |

To estimate the surface air temperature at these sites, we used Daymet, a publicly available database that estimates the surface climatology variables by interpolating the ground-based observation through statistical modeling approaches [31]. Specifically, we subset the estimated daily maximum temperature and daily minimum temperature in the year 2022 from the Daymet database. The Daymet climate data was downloaded via the programmatic interface provided by the R package *daymetr* [37]. We obtained the US state map from the US Census Bureau using the R package *tigris* [38]. We made the map in Fig. 1 with a custom R script that handles simple features using *sf* [39] and *stars* [40], and plotted the map using ggplot2 [41] and *ggspatial* [42].

### De novo genome assembly and taxonomy identification

A detailed description of genome assembly, annotation, and replicon identification are included in SI Section 2. In brief, to prepare genomic DNA for sequencing of *Sinorhizobium* genomes, we obtained cell pellets by streaking glycerol stocks on solid TY agar plates and incubated them at 30℃ for three days, followed by suspension in PBS and centrifugation. The washed pellets were resuspended in 500 μL of DNA/RNA shield for genomic DNA extraction and Oxford Nanopore long-read sequencing (Plasmidsaurus, Eugene, OR, USA). Raw reads were filtered using filtlong (ver. 0.2.1) [43] to remove the lowest 5% of quality reads, resulting in 38 samples with a median of 101 k reads, median read length of 2.7-5.4 kbp, and coverage estimates ranging from 48-192×. Downsampled reads were assembled into draft genomes using miniasm (ver. 0.3) [44]. With this draft genome, we re-downsampled the reads to a mean coverage of 100× or skipped this step if the realized coverage falls below 100×. We implemented a second downsampling using filtlong (ver. 0.2.1; options: –target_bases 100×G_d_ –mean_q_weigth 10) [43]. From these downsampled reads, we then assembled the genomes using the assembler Flye (ver. 2.9.2; options: –meta –nano-corr) [45] and polished with medaka (ver. 1.8.0) [46], achieving N50 values between 0.41-5.42 Mbp and BUSCO (ver. 5.7.1; options: -m genome -l alphaproteobacteria_odb10 -c 10) [47] completeness scores from 94.2-98.8%, indicating high assembly quality.

We determined the taxonomy via two BLAST comparisons: the first against a generic 16S rRNA gene database and the second against a custom database composed of 19 strains belonging to nine *Sinorhizobium* species, including symbiotic (*S. meliloti, S. medicae, S. fredii)* and nonsymbiotic clades (*S. adhaerens, S. mexicanum, S. canadensis, S. americanum, S. sojae*, and *S. alkalisoli*) (Table S1). We then determined the replicon identity by first removing contigs < 1 kbp and choosing the top blast hit against the custom *Sinorhizobium* database at the contig level. The contigs were categorized into four classes: chromosome, pSymA, pSymB, or pAcce according to the reference genome annotation.

### Annotation, pangenome, and phylogenomic analysis

We annotated the de novo genome assemblies using Prokka (ver. 1.14.5; options: -kingdom bacteria and --gcode 11) [48] and performed pangenome analysis using Panaroo [49]. The results included a set of individual core-gene sequence alignments (based on MAFFT [50], default in Panaroo) and gene presence-absence matrix that were both used in the phylogenetics analysis. We inferred the maximum likelihood phylogenies based on the single-copy core-gene alignments. We used a custom python script to concatenate the single-copy core-gene alignments into a single FASTA file. We then computed a maximum likelihood phylogeny using IQ-TREE2 (ver. 2.3.0; options: -nt AUTO -B 1000) with 1000 bootstrap replicates [51]. We inferred the hierarchical clustering of accessory gene content using the R function *hclust* with distance “ward.D2” based on the gene presence/absence matrix. We evaluated the sampling effort of the pangenome by fitting Heap’s law, N = k n^β^, where N is the number of orthologous genes, k is a scaling factor, n is the number of sample genomes, and β is the openness parameter.

### Growth assays

We measured the growth curves of a subset of 32 rhizobia strains at four temperatures (25℃, 30℃, 35℃, and 40℃) in the TY liquid medium. Three days before the experiments, we prepared the rhizobia inocula using a sterilized inoculation loop to streak the glycerol stock on solid TY agar plates, which were incubated at 30℃ for three days until visible colonies appeared. On the day of the experiment, we suspended each colony in a fresh TY medium and diluted the inoculum to a fixed optical density at OD_600_ = 0.1. We inoculated 2 μL of inocula to each well of a 400 μL 96-well flat-bottom clear microplate containing 100 μL of TY medium. Each isolate was assayed in two replicates at each temperature. We measured the OD_600_ using a microplate reader (BioTek Epoch 2 Microplate Spectrophotometer) every five minutes for 48 hr with incubation set at a fixed temperature without shaking. We used a custom script to fit each empirical growth curve to a generalized additive model and extracted the growth rate, lag time, and total yield for each temperature (SI Section 3).

### Statistical analysis

Unless stated otherwise, we conducted all analyses and created figures using R packages *tidyverse* [52], *janitor* [53], *cowplot* [54], and *ggh4x* [55] in R 4.4.2 [56] with deviation coding (“contr.sum”) for categorical variables. The R environment and package version were stored in a *renv* lockfile for reproducibility [57]. We performed a chi-square test to evaluate whether the two geographic regions (PA and VA) differ in the *Sinorhizobium* species isolated. We used nonlinear least squares to fit the pangenome openness using *nls* from base R. To identify the importance of thermal tolerance genes contributing to the differentiation between the two symbiotic species, we used a random forest model from the R package *randomForest* [58].

For the rest of the analysis, we either performed linear mixed models using *lmer* from the R package *lme4* [59], or in the case of nodule count, we used generalized linear mixed models using *glmmTMB* with a zero-inflated Poisson response distribution [60]. For each analysis reported in the Results below, we indicate whether a linear model (LM), linear mixed model (LMM), or generalized linear (mixed) model (GL(M)M) was used. To test whether the daily maximum temperature contributes to rhizobial growth performance, we fitted linear mixed models with a growth trait (growth rate, lag time, or yield) as the response variable, the temperature at which the growth was measured, the highest daily maximum temperature, the interaction between both, and species identity as fixed effects, as well as strains and sampling sites as random effects. The fixed effects are then tested using Type III Wald chi-square test using *Anova* from the package *car* to evaluate whether its effect differs from zero [61]. We performed post-hoc Tukey tests using the R package *emmeans* [62].

## RESULTS

### *Sinorhizobium meliloti* is more common in Pennsylvania; *S. medicae* dominates in Virginia

We first compared the daily temperature at our 15 sampling sites during the sampling season, July to September of 2022 (SI section 1) (Fig. S1). On average, the daily maximum at Pennsylvania sites is 4.8°C higher than Virginia sites (LM: χ^2^=32.6, df=1, P<0.001) (Fig. 1B). The pattern holds for daily minimum temperature (4.6°C difference; χ^2^=348.8, df=1, P<0.001). Hereafter we deemed the Pennsylvania sites as the warmer region and the Virginia sites as the cooler region (the lower latitude side was cooler because it is at higher elevation).

The de novo genome assemblies of 38 strains isolated from *M. lupulina* plants exhibit high-quality assembly with a median genome size of 7.18 Mbp (SI section 2). Analysis of the 16S rRNA gene revealed two main clades: 35 symbiotic strains and three nonsymbiotic strains, both belonging to the genus *Sinorhizobium* (Table 1) [27]. Further classification of chromosome using a custom set of reference genomes grouped the strains into four species: the symbiotic clade consisting of 22 *S. meliloti* and 13 *S. medicae* strains, and the nonsymbiotic clade including one *S. adherens* and two *S. canadensis* strains (Tables 1 and S1). To confirm the symbiosis phenotype (nodulation or not) indicated by genomes, we inoculated representative strains of each species onto the host *M. lupulina*. We found a significantly small number of nodules and reduced biomass in plants inoculated with nonsymbiotic strains, compared to the symbiotic strains (GLMM: χ^2^=20.02, P<0.001 for nodules; LMM: χ^2^=11.56, P<0.001 for biomass) (Fig. S2). No significant difference was detected between *S. medicae-* and *S. meliloti*-inoculated plants in terms of nodule number (GLMM: χ^2^=0.0014, P=0.97) or biomass (LMM: χ^2^=0.96, P=0.33).

The two symbiotic species show regional dominance: *S. meliloti* is more frequently found in the warmer Pennsylvania sampling sites, whereas *S. medicae* predominates in the cooler Virginia area (Fig. 1C) (chi-square test: χ^2^ =20.73, df=3, P<0.001). All three nonsymbiotic strains were isolated exclusively from the cooler sampling sites in Virginia.

### Thermal tolerance differed between *Sinorhizobium* species

We hypothesized that *Sinorhizobium* strains exhibit thermal local adaptation, with the dominant species from the warmer sampling sites (*S. meliloti*) exhibiting greater heat tolerance compared to dominant species from the cooler sampling sites (*S. medicae*). To test this, we assessed their heat tolerance as growth in liquid culture across four temperature settings—25°C, 30°C, 35°C, and 40°C— spanning the upper range of daily maximum temperatures at our sampling sites (Fig. 1B). Our results reveal species-specific thermal responses (Fig. 2A). At 25°C and 30°C, nonsymbiotic strains exhibited higher growth rates compared to symbiotic strains (LMM: χ^2^=17.68, P<0.001 for 25°C and χ^2^=7.38, P=0.006 for 30°C). This difference disappeared at the higher temperatures (LMM: χ^2^=0.42, P=0.52 for 35°C and χ^2^=0.73, P=0.39 for 40°C) (Fig. 2B). Notably, at 40°C, *S. medicae* and nonsymbiotic strains showed minimal growth after 48 hours, indicating intolerance to high temperature. By contrast, *S. meliloti* maintained growth at 40°C at a reduced or similar level compared to its growth at other three temperatures (Fig. 2B) (post hoc Tukey with regard to temperature). Between the two symbiotic species, *S. meliloti* generally has 0.08hr^-1^ higher growth rates, 6.1 hr shorter lag phase, and 0.14 higher yield than *S. medicae* throughout the tested temperatures (post hoc Tukey with regard to species). This difference is nuanced at the lower temperatures and becomes pronounced at 40°C where *S. meliloti* achieved significantly higher growth due to *S. medicae* intolerance to the heat (LM: t=2.28, P=0.03 for growth rate; t=5.06, P<0.001 for lag time; t=7.35, P<0.001 for yield). These findings are consistent with the observed biogeographic pattern of *Sinorhizobium: S. meliloti* was more common in the warmer sites, while *S. medicae* was more common in the cooler sites.

**Figure 2.**
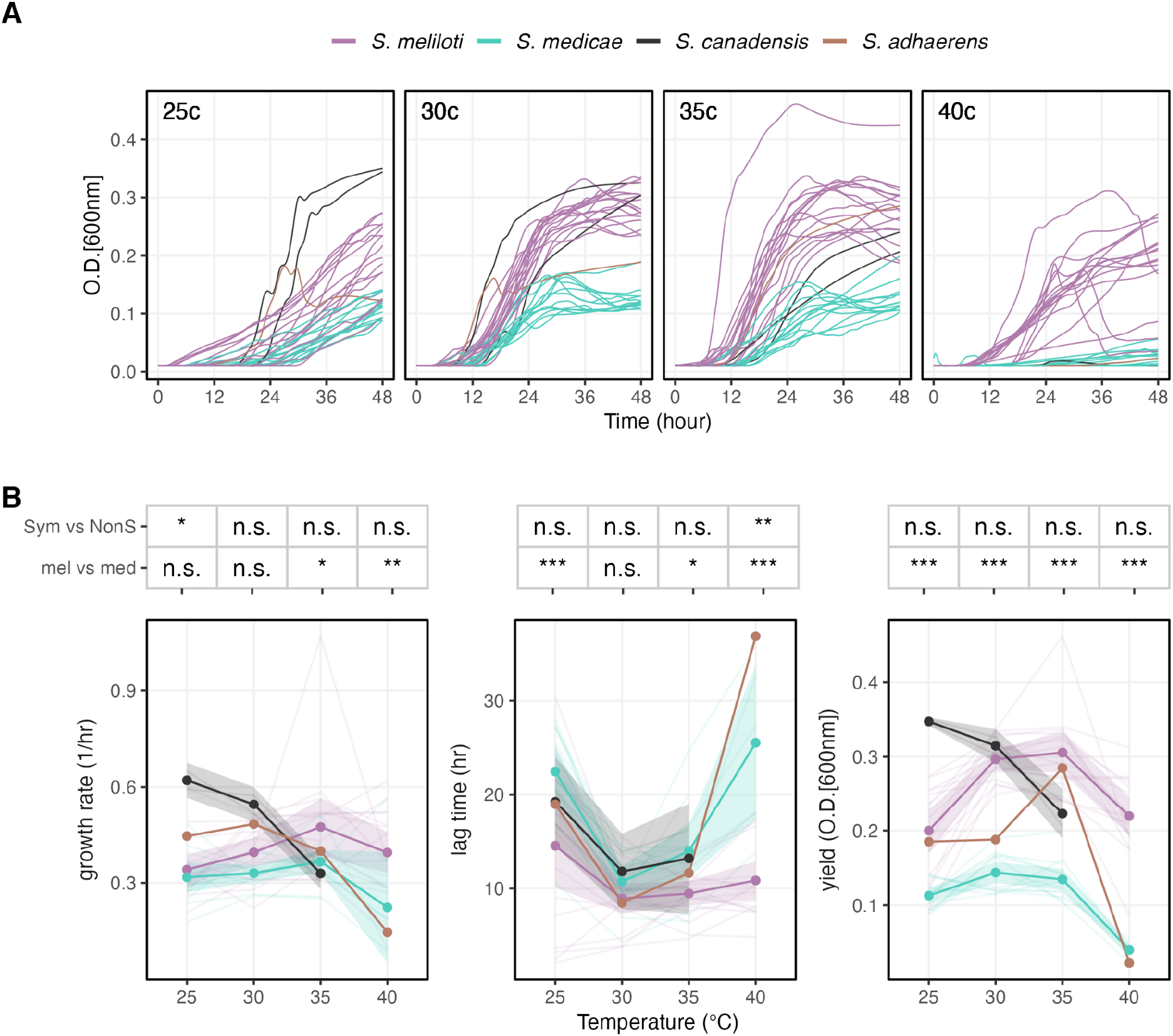
Free-living growth traits in four different temperatures. (A) We measured the growth curve of free-living rhizobia at 5-min intervals for 48hr on a nutritious liquid medium. Each line represents the smoothed, average optical density of two replicates per strain. (B) The semitransparent lines represent the strains whereas the solid dots denote the mean values and shades represent the 95% confidence intervals. *S. adhaerens* only has one strain so no shade is shown. Note that 10 strains have at least one replicate that did not grow over OD = 0.01 at 40°C, so estimating the growth traits under such conditions is not possible. The asterisks above each temperature represent the statistical significance ( P<0.001***, P < 0.01**, P<0.05*, or nonsignificant n.s) of post hoc Tukey test between symbiotic and nonsymbiotic strains (“Sym vs NonS”) and between *S. meliloti* and *S. medicae* (“mel vs med”).

We then asked if the temperature features of our sampling sites are predictive of rhizobia growth traits. If the daily maximum temperature drives the thermal adaptation in rhizobia, we should expect it to be correlated with growth performance. After accounting for the contribution of species difference, we found that daily maximum temperature significantly contributed to the variation in strains’ lag time (LMM: χ^2^=5.58, P=0.018) and yield (LMM: χ^2^=5.06, P=0.024), but not the growth rate (LMM: χ^2^=0.028, P=0.87). However, the effect of daily maximum temperature is gone within regions, except for the yields in the warm region (LMM: χ^2^=5.16, P=0.023). Together, these results suggest that the rhizobial thermal adaptation occurs at regional scale, but not as pronounced among sites within a region.

### Contrasting genomic features of *S. medicae and S. meliloti*

To investigate the population structure of *Sinorhizobium* within and between regions, we compared 38 de novo genome assemblies. Pangenome analysis identified 987 core genes, accounting for approximately 3.72% of the total 26,544 genes (core/total=23.3% of 16,166 genes for symbiotic clade and core/total=24.8% of 11,509 genes for nonsymbiotic clade) (Fig. S3). Phylogenetic analysis based on both 775 single-copy core genes and hierarchical clustering of accessory gene content revealed three major clades (Fig. 3AB) (Fig. S4), consistent with taxonomic assignments based on chromosome identity (Table 1). Tree distance analysis also suggests a congruence between core and accessory genomes (Fig. S4).

**Figure 3.**
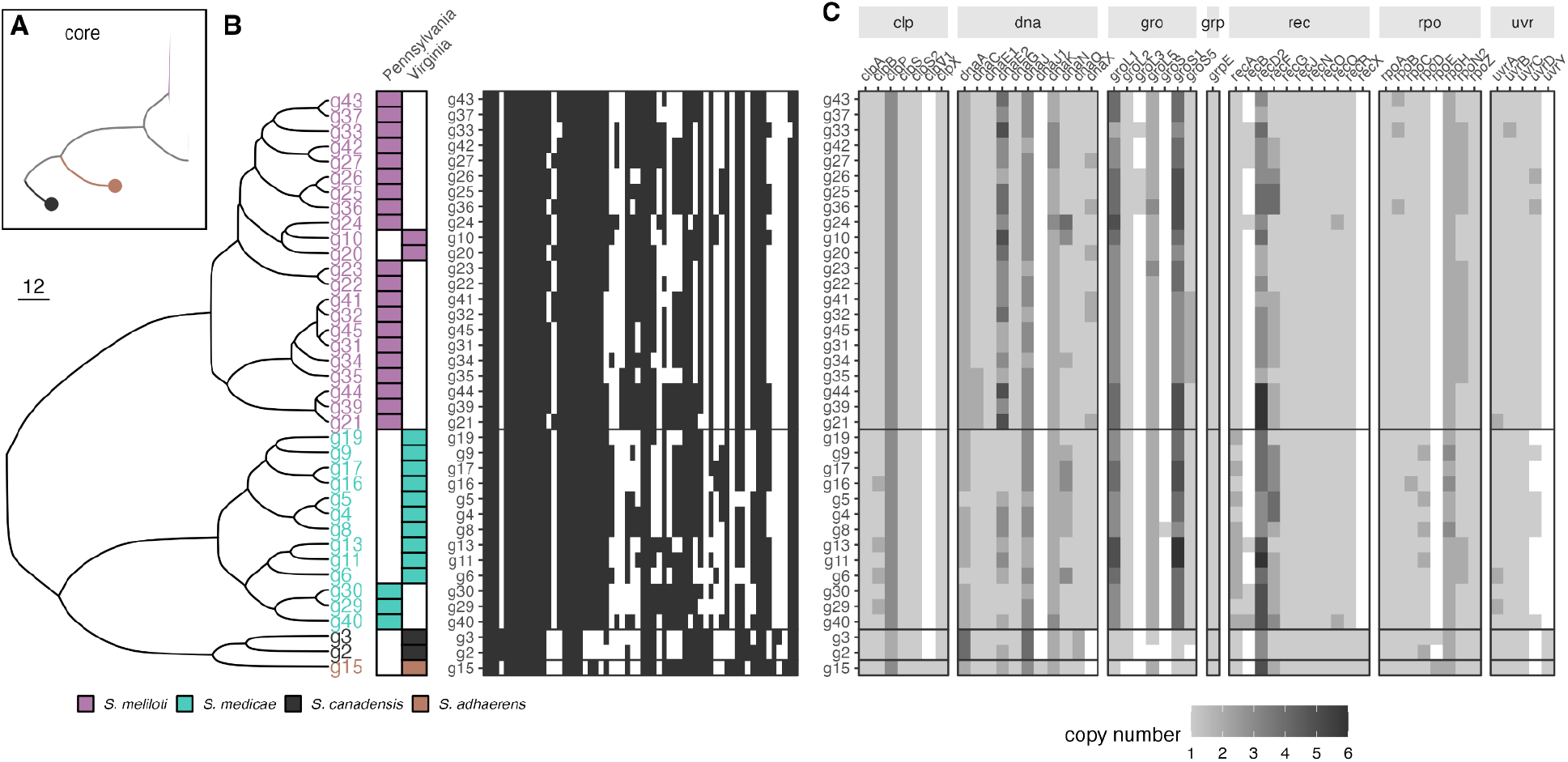
Phylogeny and gene content variation. (A) Phylogeny based on 775 single-copy core-genes. Conspecific strains are collapsed into single clades. All internal nodes have > 99.9% bootstrap support. (B) Hierarchical clustering of *Sinorhizobium* strains based on the heat map of 26,544 genes. The region of origin is indicated in the center. Presence and presence of the homolog for each gene are indicated in black and white, respectively. (C) 38 genes relevant to heat tolerance.

We compared the gene content shared or specific to each symbiotic species. We excluded the nonsymbiotic species from this analysis because of low sample size. Within the two symbiotic species (*S. meliloti* and *S. medicae*), the proportions of core and accessory genes are similar when accounting for the number of genomes analyzed (core/total=45.9% for *S. meliloti* and 45.3% for *S. medicae*) (Fig. S5). The openness parameter (β) according to Heap’s law is 0.16 for *S. meliloti* and 0.19 for *S. medicae*, suggesting a fairly closed pangenome in both symbiotic species (Fig. S5).

We then compared accessory genome content to determine whether there were differences in the proportion of the genome attributed to heat-relevant genes. We found that the two symbiotic species differ in the composition of the 38 functional genes relevant to heat tolerance (PERMANOVA: F=9.56, P=0.001) (Fig. 3C). Among these functional genes, we found that the composition of recA, uvrD, and dnaE2 together explains 44.5% of the variance, with recA enriched in *S. medicae* and uvrD and dnaE2 enriched in *S. meliloti* (Fig. S6), suggesting a possible target of selection underlying the differentiation in thermal tolerance between the two symbiotic species.

### Gene flow is more restricted among the Pennsylvania sites than the Virginia sites

Finally, we investigated the local population structure of *Sinorhizobium* within regions. Strains from the same sampling sites tended to cluster together on both the single-copy core-gene phylogeny and the hierarchical clustering of accessory genome content (Fig. S4), suggesting limited gene flow within each region. To test this, we examined patterns of isolation by distance within each geographic region for each replicon (Fig. 5). We focused on the most locally abundant species in each region (*S. meliloti* in Pennsylvania and *S. medicae* in Virginia) to minimize interspecific variation. We found significant isolation by distance for the *S. meliloti* chromosome, pSymA, and pSymB (Fig. 4A). A similar pattern holds for gene content variation (Fig. 4C). In contrast, for *S. medicae*, we did not detect significant isolation-by-distance for any replicon, either based on core gene sequence similarity or accessory genome content (Fig. 4BD). These results suggest that gene flow might homogenize the populations of *S. medicae* in Virginia, but is more limited in *S. meliloti* in Pennsylvania.

**Figure 4.**
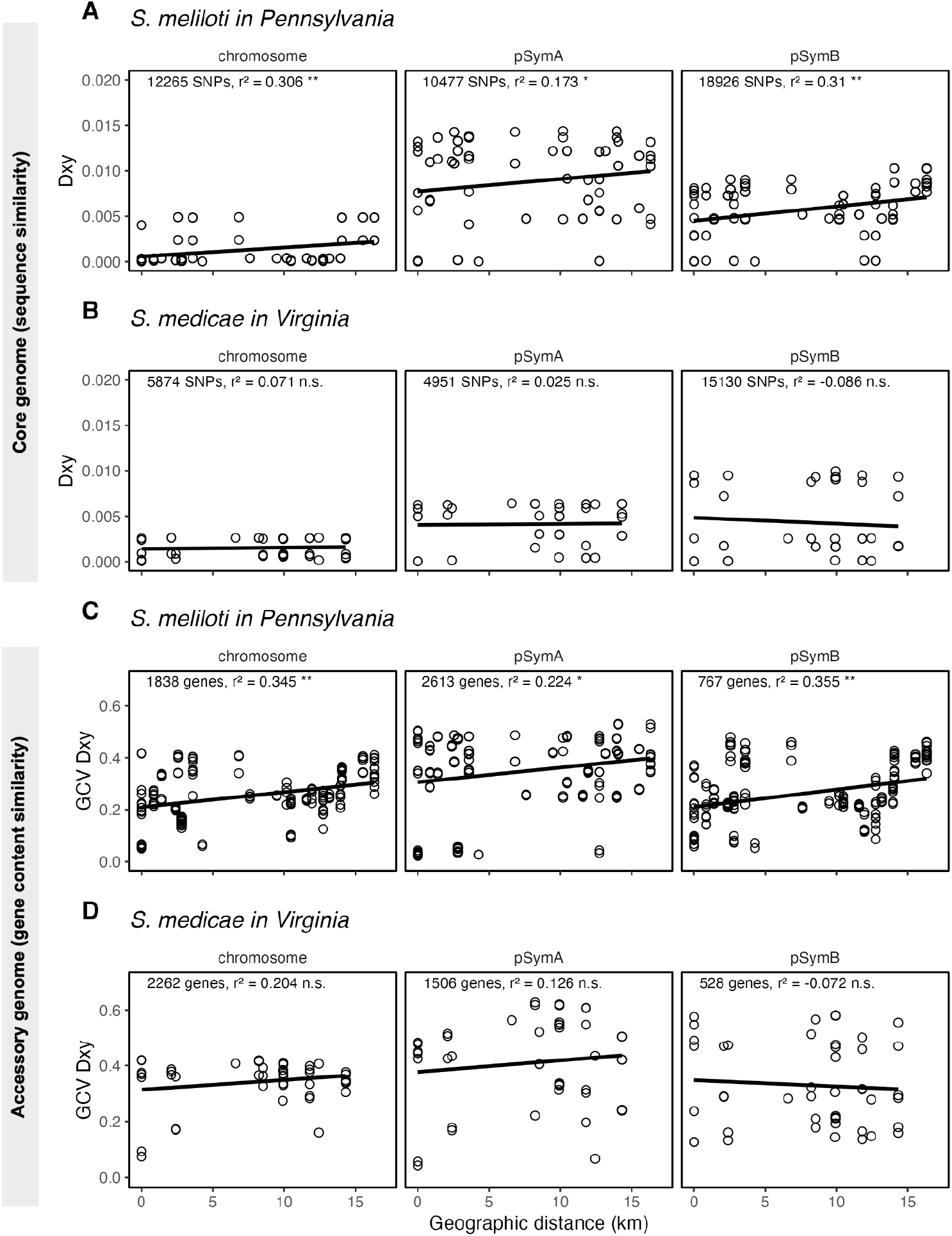
Gene flow is more restricted for *S. meliloti* among the Pennsylvania sites. (A) and (B) show the nucleotide distance of single-copy core genes between two rhizobia genomes. (C) and (D) show the Manhattan distance (scaled by number of accessory genes) of gene content between two genomes. Mantel tests are reported in each panel shown with the number of loci (SNPs or accessory genes), and statistical significance (n.s. non-significant, * p< 0.05, ** p<0.01). Note that the geographic distance spanned by both gradients is nearly identical (∼15 km in both cases).

## DISCUSSION

Although facultative symbiosis is a common strategy across the tree of life, our understanding of how adaptation in its free-living stage contributes to the biogeography of facultative symbionts is still growing. In this study, we investigated the biogeographic distribution and thermal local adaptation of its facultative endosymbiont, *Sinorhizobium*. Laboratory growth assays revealed variability in thermal tolerance among species: *S. meliloti* maintained growth at high temperatures, while *S. medicae* and non-symbiotic species failed to grow under similar conditions. These differences align with the observed biogeographic patterns of these rhizobia. Population genomics indicated an isolation-by-distance pattern in the most prevalent symbiotic species within a particular region, suggesting that limited gene flow, rather than thermal adaptation, operates at a finer spatial scale.

### *Sinorhizobium* exhibit thermal local adaptation

Our growth assays demonstrated that *S. meliloti* exhibits greater heat tolerance than *S. medicae* and non-symbiotic species, aligning with the thermal environments from which these strains were isolated. Our growth assays also demonstrate that among *M. lupulina*-nodulating rhizobia, *S. meliloti* exhibits a higher degree of thermal tolerance, maintaining growth at 40°C, whereas *S. medicae* maximum growth temperature falls between 35°C and 40°C. These findings align with climatological data from our sampling sites, which show that the *S. medicae*-dominated sites generally experience maximum temperatures below 35°C, while the *S. meliloti*-dominated sites frequently exceed this threshold. The biogeographic patterns in our data also support a previous study examining the population structure of *M. lupulina* across eastern North America, which found *S. medicae* to be more prevalent in cooler, northern regions and *S. meliloti* more common in the warmer south [35]. A previous study tested the hypothesis that this pattern is attributable to rhizobial adaptation to local hosts, but reciprocal inoculation experiments found no supporting evidence [63]. Our growth assays suggest an alternative hypothesis: the regional geographic distribution of these symbionts may be driven by adaptation to non-host environmental factors such as thermal tolerance.

Historical studies on temperature effects on rhizobia growth corroborate these results. Allison et al (1940) examined nine strains across five rhizobium species, including *R. meliloti* (the cryptic species equivalent to *S. meliloti*), and found *R. meliloti* to be the most heat-resistant, with an optimal temperature range of 28–30°C, tolerating up to 39°C, but losing growth at 41°C [26]. Similarly, Bowden and Kennedy (1959) tested several rhizobium strains from legumes including *Medicago sativa* (alfalfa), reported maximum temperature tolerances ranging from 36.5°C to 42.5°C [64]. Additionally, soil temperature has been shown to strongly influence nodulation rates in legumes; for example, studies on beans have shown that optimal growth occurs at around 25°C rather than 30°C, even under equal inoculation conditions, highlighting the complex relationship between temperature and nodulation efficiency [65]. These pioneering studies on rhizobia thermal performance highlight the effectiveness of simple growth assays in offering deep insights into how temperature influences rhizobial distribution and function across their two life stages in diverse environments.

Our comparative genomics analysis identified three accessory genes—recA, uvrD, and dnaE2—as potential factors underpinning the differences in thermal tolerance between *S. meliloti* and *S. medicae*. uvrD and dnaE2 were more abundant in *S. meliloti*, consistent with the idea that enhanced DNA repair and stress-induced mutagenesis facilitate adaptation to elevated temperatures. uvrD, a gene involved in DNA repair, likely contributes to higher thermal tolerance by increasing repair efficiency at elevated temperatures and also plays a role in desiccation resistance [66]. dnaE2, associated with stress-induced mutagenesis, is present in multiple copies in *S. meliloti*, potentially aiding high-temperature adaptation by facilitating the evolvability of stress-induced mutagenesis, therefore adaptation under thermal stress [67]. By contrast, recA was relatively enriched in *S. medicae*. recA is essential for homologous recombination and the repair of double-stranded breaks, yet the reduced representation of recA in *S. meliloti* suggests that high thermal tolerance can evolve through mechanisms that partially bypass canonical homologous recombination repair [68]. One possibility is that *S. meliloti* relies more heavily on uvrD and dnaE2–mediated repair and mutagenesis pathways under thermal stress, decoupling thermal adaptation from recA-mediated recombination. Notably, recA mutants do not impact its symbiotic nitrogen fixation, suggesting that thermal adaptation and symbiosis can be decoupled via recA variation [69]. Together, the presence, variation, or regulation of these genes may explain *S. meliloti’s* capacity to withstand higher temperatures, providing insights into the genetic mechanisms of thermal adaptation in these symbiotic bacteria.

Our findings add to this growing body of evidence highlighting abiotic factors, specifically thermal tolerance, as important agents of selection in free-living rhizobia, and underscore the importance of characterizing off-host adaptation to understand the biogeography of facultative symbionts. Support for this hypothesis also comes from studies on rhizobia adaptation to soil conditions. Among soybean-nodulating rhizobia, *Sinorhizobium* is widespread in alkaline-saline soils, while *Bradyrhizobium* dominates in neutral to acidic soils [13]. Additionally, in heavy metal-rich serpentine soils, *Mesorhizobium* have shown increased frequencies of alleles and genes that confer heavy metal tolerance [12].

### Isolation by distance varies across *Sinorhizobium* species and regions

In *S. meliloti* populations from Pennsylvania, we detected isolation by distance (IBD) in both core gene SNPs and accessory gene content, consistent with our phylogenomic congruence test and indicating concordant population structure across genome compartments. The strength of IBD differed across species and regions: *S. meliloti* in Pennsylvania exhibited stronger spatial structure than *S. medicae* in Virginia. This contrast suggests that the potential for population divergence may be shaped by both species-specific traits and landscape context. One one hand, intrinsic differences in dispersal ability or recombination rates could contribute to the observed variation [70, 71]. Alternatively, regional environments may impose distinct constraints on dispersal opportunities [15, 72, 73]. For instance, the Pennsylvania sites were primarily urban, where microbial populations may experience restricted local dispersal due to habitat fragmentation, whereas the more continuous habitats in Virginia may facilitate greater connectivity. At the same time, urbanization has been shown to homogenize microbial communities across sites by reducing regional spatial turnover [74, 75]. These studies along with our results highlight how urban environments may simultaneously constrain local microbial movement while promoting broader-scale similarity, producing distinct IBD patterns in urban populations [76]. Taken together, disentangling species-level traits from regional influences will require broader sampling across multiple regions to separate the contributions of intrinsic biology and environmental context to microbial population structure.

## Conclusion

This study examined how evolution during the free-living phase influences the biogeographic patterns of *Sinorhizobium* rhizobia, a well-known facultative symbiont associated with legume plants. By integrating laboratory growth assays with targeted sampling, the research demonstrates the effectiveness of simple, trait-based methods in elucidating the biogeography of facultative symbionts, complemented by comparative genomics to identify potential genomic targets of selection. While this approach is limited to culturable strains, we anticipate that future studies will place greater emphasis on the free-living stages of facultative symbionts, thereby advancing our understanding of the evolutionary and ecological factors that contribute to the stability of microbes with dual life histories.

## ACKNOWLEDGEMENTS

We thank the members of the Wood lab for their feedback on this manuscript. We would like to thank the staff at Mountain Lake Biological Station for their support on the fieldwork. Growth chamber logistical support was provided by the University of Pennsylvania’s Greenhouse support staff members Kathyrn Butler and Samara Gray. This project was supported by NSF-DEB 2118397 awarded to C.W.W, and startup funding from the University of Pennsylvania. C.-Y. C. was supported by The Data-Driven Discovery Postdoctoral fellowship by the University of Pennsylvania. Gerardo Bencosme was supported by the Mountain Lake REU program.

## Statement of Authorship

C.-Y.C. and C.W.W conceived the idea and designed the study. G. B. performed the preliminary experiments. C.-Y.C., and T.T.-B., and J.L.R. performed the experiments and collected the data. C.-Y.C., M.B.C., and C.W.W discussed the results and drafted the paper. C.-Y.C. and C.W.W. wrote the final version of the paper.

## Data and Code Availability

Scripts and curated dataset needed to replicate results, figures, and tables in this article are available on Zenodo https://doi.org/10.5281/zenodo.16986125. All scripts used to process from raw to final presentation objects are also included, whereas the raw sequences and intermediate data will be accessible upon acceptance for the sake of storage limit.

