## Supplementary material for "Biogeographic and genomic signatures of thermal adaptation in facultative rhizobia": SI

**Supplementary information for the manuscript entitled "Biogeographic and genomic signatures of thermal adaptation in facultative rhizobia"**

This includes:
Supplementary Sections 1-4
Supplementary Figures 1-6
Supplementary Table 1

#### Supplementary Section 1: Sampling Sites

To estimate the surface air temperature at these sites during the field season, we used Daymet, a publicly available database that estimates the surface climatology variables by interpolating the ground-based observation through statistical modeling approaches (Thornton et al. 2022). Specifically, we subset the estimated daily maximum temperature and daily minimum temperature in the year 2022 from the Daymet database. The Daymet climate data was downloaded via the programmatic interface provided by the R package *daymetr* (Hufkens et al. 2018).

*Maps.* We obtained the US state map from the US Census Bureau using the R package *tigris* (Walker 2024), the elevation data using *geodata* (Hijmans et al. 2023), and year 2022 fraction impervious surfaces using the Multi-Resolution Land Characteristics Consortium Viewer (https://www.mrlc.gov/viewer/). Specifically, from MRLC we selected a rectangular area ranging between longitude [75.5°W, 75.1°W] and latitude [39.7°N, 40.1°N]. We made the map in Fig. 1 with a custom R script, where we computed the distance between sites using *geosphere* (Hijmans 2024), handling the simple features using *sf* (Pebesma 2018) and *stars* (Pebesma and Bivand 2023), and plotted the map using ggplot2 (Wickham 2016) and *ggspatial* (Dunnington 2023).

#### Supplementary Section 2: Genome assembly and annotation

We used bioconda/miniconda (Grüning et al. 2018) and mamba to manage the interdependency of the bioinformatic tools. In principle, we created one mamba environment for each tool mentioned below to avoid conflicts in package dependency. The shell script *setup_conda.sh* sets up miniconda on macOS whereas *setup_envs.sh* creates the mamba environments. The shell scripts were executed on a 2021 iMac with Apple M1 chip 16GB memory and macOS version 14.7.

##### ***De novo*** **assemblies of** ***Sinorhizobium*** **genomes**

*Rhizobia cell pellets.* To prepare the cell pellets for genomic DNA extraction and sequencing, we streaked the glycerol stocks onto solid TY agar petri plates and incubated the streaked plates at 30℃ for five days. We then suspended the colonies on a plate using sterile pipette tips in 1 mL of PBS, followed by centrifugation at 14000 rpm for 2 mins and the removal of supernatants. We then re-suspended the pellets in 0.5 mL of DNA/RNA shield (Zymo Research) and sent the cell samples for the "Big with Extraction" genomic DNA extraction and Oxford Nanopore long-read sequencing service provided by Plasmidsaurus, Eugene, OR, USA.

*Raw read quality control.* Following the *de novo* assembly workflow recommended by Plasmidsaurus, we first use filtlong (ver. 0.2.1) (Wick and Menzel 2017) to remove the raw read with the worst 5% quality in each sample. Among the 38 filtered samples, the number of raw reads per sample ranges from 47-234 kbp with a median of 101 kbp. Across the 38 samples, the median read length ranges from 2.7-5.4 kbp and the mean read quality score ranges from 18.26-22.54. The estimated read coverage across the 38 samples ranges from 48-192$\times$, assuming the target genome size of the representative *Sinorhizobium meliloti* at 7 Mbp (Galibert et al. 2001). The sequence coverage allowed us to perform reference-free *de novo* assembly.

*De novo assembly*. To reduce computational costs, we downsampled the filtered reads to a total of 250 Mbp using filtlong (ver. 0.2.1) (Wick and Menzel 2017) by removing reads with worse quality. We then assembled a draft genome of size $G_d$ using miniasm (ver. 0.3) (Li 2016). With this draft genome, we re-downsampled the reads to a mean coverage of 100$\times$ or skipped this step if the realized coverage falls below 100$\times$. We implemented a second downsampling using filtlong (ver. 0.2.1; options: –target_bases 100$\times G_d$ –mean_q_weigth 10) (Wick and Menzel 2017). From these downsampled reads, we then assembled the genomes using the assembler Flye (ver. 2.9.2; options: –meta –nano-corr) (Kolmogorov et al. 2019). We polished the consensus genomes using medaka (ver. 1.8.0) (*Medaka: Sequence Correction Provided by ONT Research* 2018) with raw read data with the model r941_min_high_g303. We then removed the contigs < 100 kbp. Finally, we manually checked the contig size and blast results (see section below) and concatenated contigs in a genome: in g20 we concatenated contigs 3, 7, 9 into one contig; in g24, we concatenated contigs 10, 17, 9, 14, 18 into one contig. The concatenation was performed using a custom shell script *manual_concat.sh*.

*Assembly quality control.* The assemblies were evaluated in two dimensions: contiguity, which evaluates both the size and quantity of contigs; and completeness, which determines the presence and absence of highly conserved genes. We assessed contiguity using QUAST (ver. 5.2.0; option: -t 10) (Gurevich et al. 2013). The N50 ranged between 0.41-5.42 Mbp, and L50 ranged between 1-7. We assessed completeness using BUSCO (ver. 5.7.1; options: -m genome -l alphaproteobacteria_odb10 -c 10) (Simão et al. 2015) with the lineage setting to alphaproteobacteria that has 432 BUSCOs. The BUSCO scores range between 94.2- 98.8%, indicating a good assembly quality. The resulting genome size ranges from 6.79-8.12 Mbp with a median of 7.18 Mbp, and the number of > 100 kbp contigs per genome is 3-5, corresponding to the previously reported genome sizes and replicon numbers of four representative *Sinorhizobium* species, *S. meliloti* (Galibert et al. 2001), *S. medicae* (Reeve et al. 2010), *S. fredii* (Weidner et al. 2012), and *S. adhaerens (Williams et al. 2017)*.

**Rhizobia taxonomy and contig identification**

We identified the genomes using two approaches that perform searches with varied specificities. First, we performed a search using only rRNA sequences from the assembled genomes. We used barrnap (ver. 0.9) (T. Seemann, n.d.) to acquire 16S rRNA sequences and blasted the rRNA sequences to a RefSeq 16S rRNA database (https://www.ncbi.nlm.nih.gov/refseq/targetedloci/16S_process/) composed of 27114 16S rRNA sequences. Second, we performed a search by blasting the assembled genomes to a custom database composed of 19 strains belonging to nine *Sinorhizobium* species (*S. meliloti, S.medicae, S. adhaerens, S. mexicanum, S. canadensis, S. fredii, S. americanum, S. sojae,* and *S. alkalisoli*) (Table S2) with BLAST (ver. 2.14.1: options: blastn -outfmt '6 qseqid sseqid pident length mismatch gapopen qstart qend sstart send evalue bitscore' -num_alignments 5). We reported the top contig hits and 16S rRNA hits for each genome in Table 1.

**Genome annotation and pangenomic analysis**

We annotated the 38 genomes with Prokka (ver. 1.14.5; options: -kingdom bacteria and --gcode 11) (Torsten Seemann 2014). Note that we excluded g42, a *S. meliloti* strain, from pangenome analysis because the genome assembly does not resolve the three main replicons. The pangenome was inferred using Panaroo (ver. 1.3.4; options: -a core --aligner mafft --core_threshold 0.95 -t 10 --clean-mode strict --remove-invalid-genes) (Tonkin-Hill et al. 2020) with the mode set to strict and with core gene alignment using MAFFT (ver. 7.520) (Katoh et al. 2002). This resulted in two datasets: multiple sequence alignment for each core gene, where single-copy core genes were later used to infer a maximum likelihood phylogeny; a gene presence-absence table indicating whether an orthologue is present in a genome. Figure S3 shows the pangenome composition and the gene frequency spectrum.

#### Supplementary Section 3: Growth assays

*Microplate reader setting*. At the later stage of incubation, uneven water evaporation may alter the liquid volume in individual microplate wells, which in turn introduces measurement errors in the optical density. To account for evaporation, the microplate reader was set to consider the path length correction set at a test wavelength of 977 nm, a reference wavelength of 900 nm, and the K-factor at 0.18. To further account for liquid volume change throughout incubation, in our plate design, we intentionally left 32 wells uninoculated as the cell-free control and blank.

*Growth traits estimation.* Following the approach described before (Estrela et al. 2022), we smoothed and extracted three growth traits from the growth curves: lag time, maximum growth rate, and maximum yield. Briefly, we first subtracted the $OD_{600}$ of cell cultures from the cell-free cultures as blank. To compute the maximum growth rate, we smoothed the $\log(OD_{600})$ of each replicate by fitting a generalized additive model using the *gam* function from the R package *mgcv* (Wood 2011). We excluded the first 1 hr and any $OD_{600}$ below 0.015 to avoid artifacts introduced by measurement and fitting noises, respectively. We then took the maximum derivative as the maximum growth rate. Lastly, the yield is determined by the maximum OD reading. As a result, we obtained 12 traits per strain (three growth traits at four temperatures) from the growth curve experiments.

#### Supplementary Section 4: Plant inoculation experiment

We chose six representative strains belonging to the four species (*S. meliloti, S. medicae, S. adhaerens*, and *S. canadensis*) and inoculated them onto 15 plant maternal lines of sterile *M. lupulina* seedlings that were collected from wild populations during the same season as rhizobia sampling. To initiate the experiment, we scarified the seeds with a razor blade, sanitized them using ethanol and household bleach, and stratified them in Petri plates wrapped in aluminum foil to block light at 4℃ for five days. After 12-16 hours at room temperature to encourage radicle elongation seedlings were planted in autoclaved cone-tainers (Stuewe & Sons, Inc., Tangent, OR, USA) filled with a 1:4 mix of turface and sand. The plants were maintained in a growth chamber with a 16:8 hour light:dark cycle, maintaining daytime temperatures at 22℃, nighttime temperatures at 18℃, and a relative humidity of approximately 50%. Five weeks after rhizobial inoculation, we top-fertilized individual plants with 5 mL of one-fifth diluted nitrogen-free Fahraeus fertilizer (Barker et al. 2006) twice a week using an electronic peristaltic pump (Integra DOSE-IT with 4mm silicone tubing) and bottom-watered the plants once a week.

One week after planting, we inoculated each plant with 1 mL of rhizobium inoculum, which had been grown in TY medium at 30℃ for two days and then diluted to an $OD_{600} = 0.1$ using fresh TY medium. To prevent cross-contamination, we arranged the cone-tainers in groups of six, with each group sharing a single water reservoir to collect excess water. These blocks were included as a random effect in our analysis. Notably, the negative control, which received only fresh TY medium, showed some nodulation, suggesting that a low level of cross-contamination may have occurred.

At the end of the sixth week, we harvested 167 plants (including 8 controls), and measured dry shoot biomass, dry root biomass, and counted the nodules. The above-ground portions were dried in a 60℃ oven, while the below-ground parts were frozen at -20℃ for up to six months prior to phenotyping. For phenotyping, we thawed each frozen root sample and counted the nodules under dissecting microscopes. The root samples were then dried at 60℃ and weighed using a scale with a resolution of 0.1 mg.

### SUPPLEMENTARY FIGURES

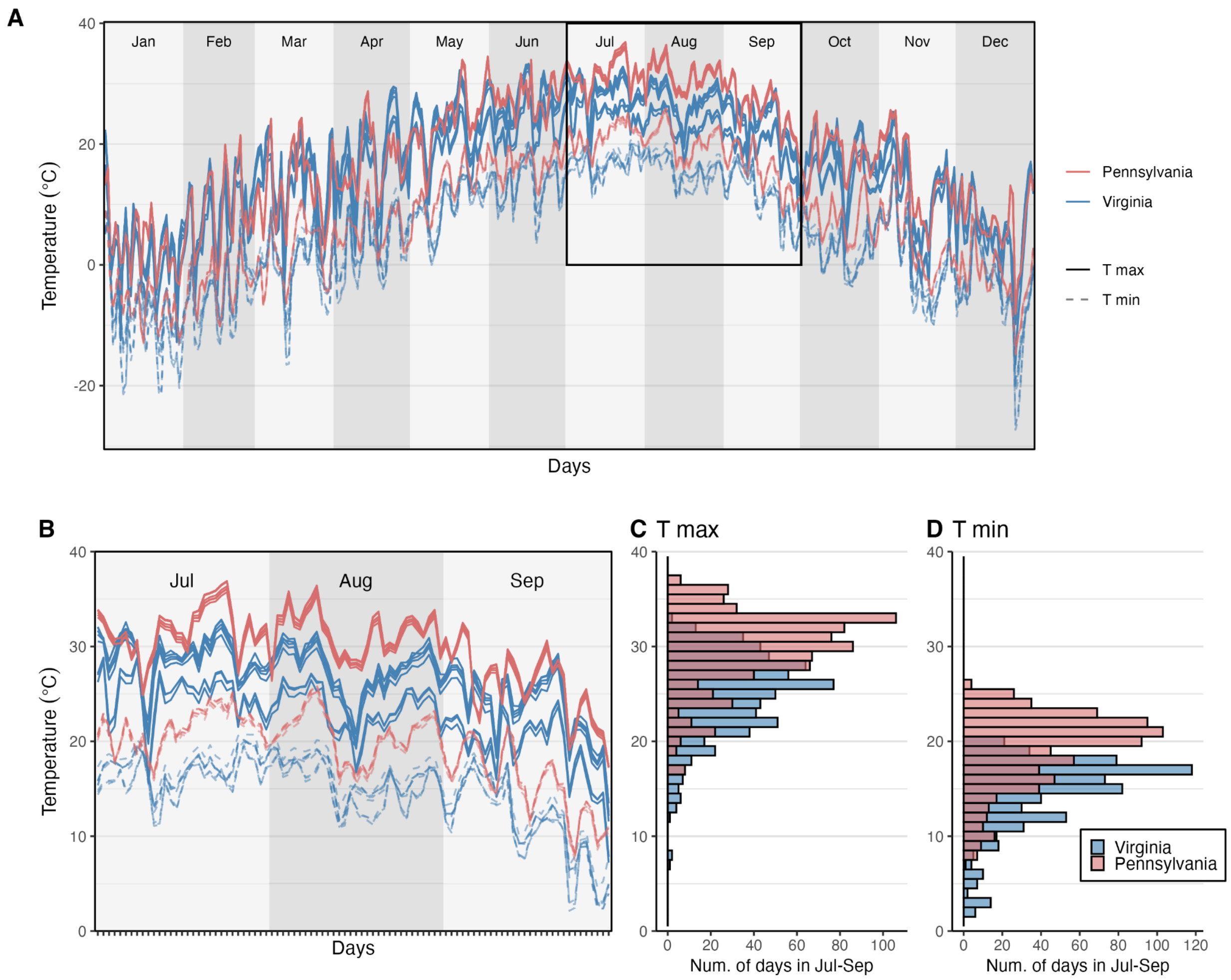


**Figure S1. Daily temperature in the regions (Pennsylvania and Virgina) of sampling.** (A) Daily temperature maximum (T max) and minimum (T min) in the year of 2022. (B) Daily temperature in the sampling season July to September. (C) Histogram of daily maximum by the number of days during July to September. (D) Daily minimum.

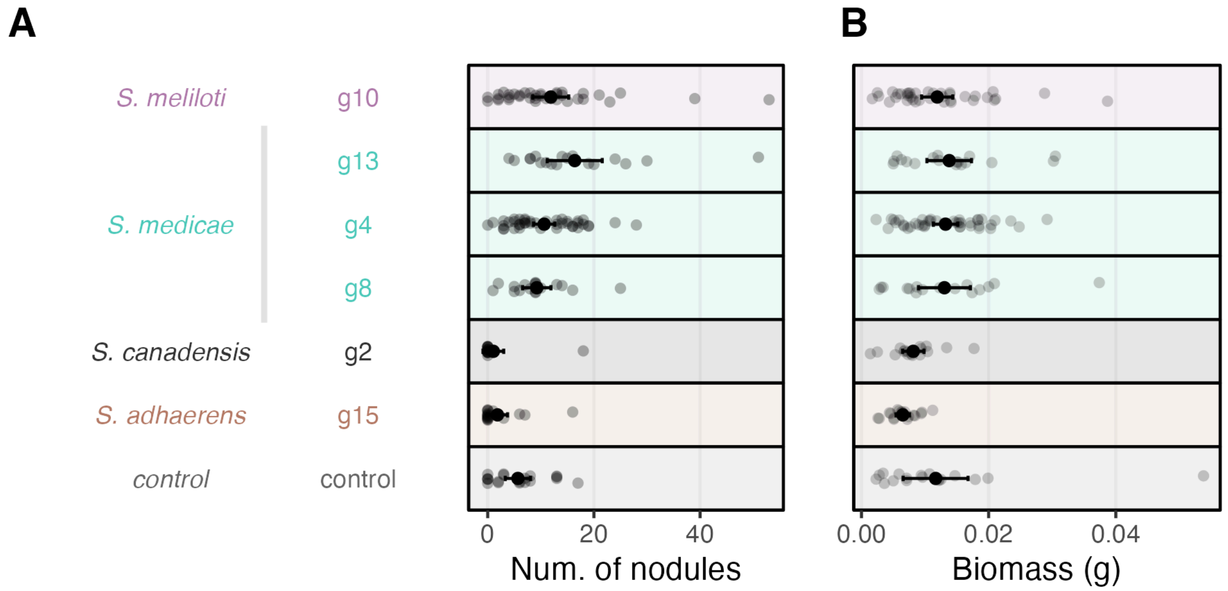


**Figure S2. Symbiosis traits of *Sinorhizobium* species.** Each transparent point presents the measurements of an inoculated *M. lupulina* plant. The solid points are the means and the error bars denote the 95% confidence intervals. (A) Number of nodules per plant. (B) The biomass of a plant = aboveground + belowground biomass.

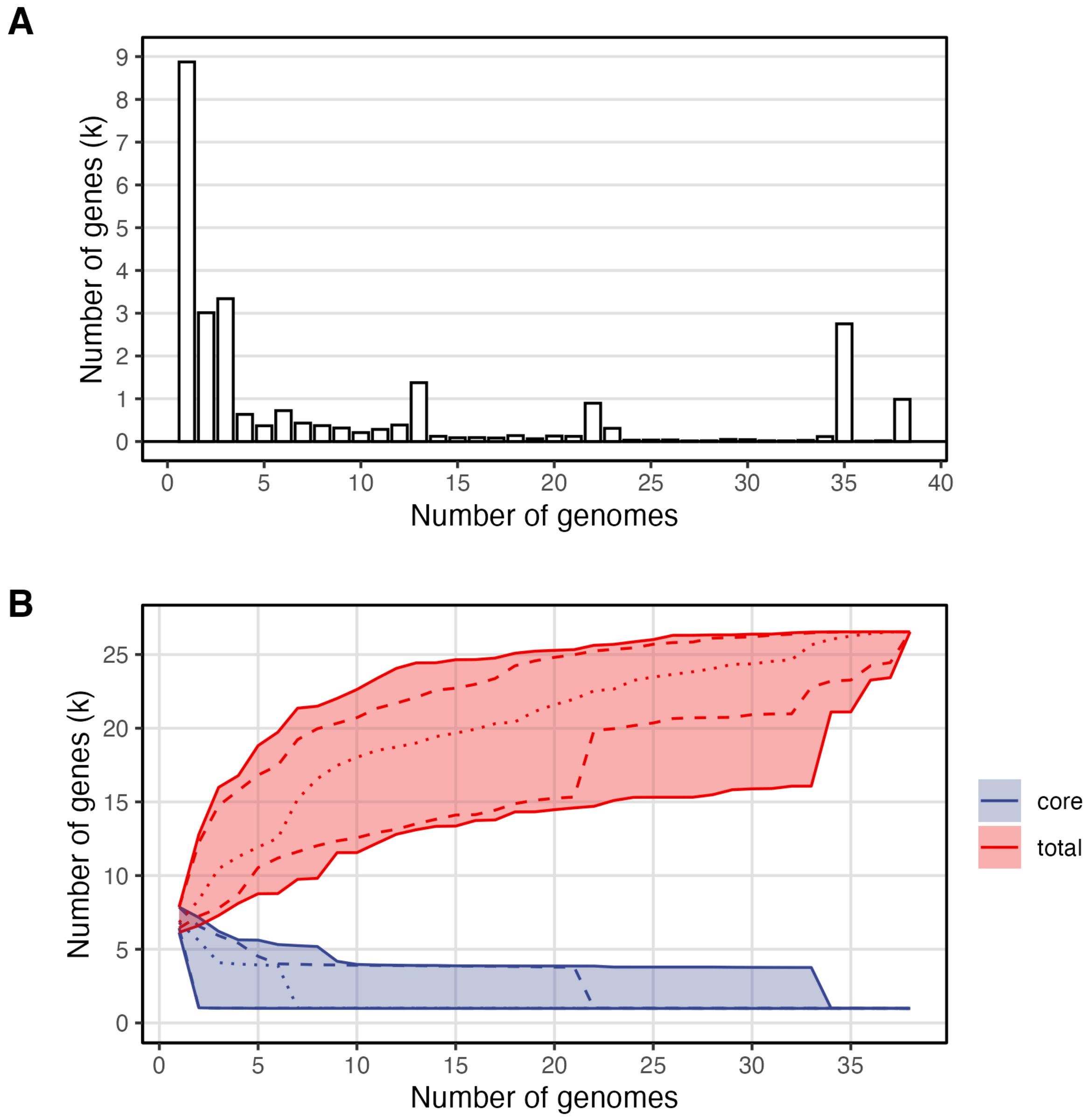


**Figure S3. Pangenome sampling.** (A) Gene frequency spectrum shows the 8875 singletons (i.e., only present in one but not other genomes), 987 core genes (i.e., shared by all genomes; including 775 single-copy core genes), and a total of 26544 genes in the pangenome of 38 *Sinorhizobium* genomes. (B) We estimated the sampling effort of genomes. At each number of genomes, we performed 100 bootstraps and computed the number of core genes and the number of total genes. The solid lines represent the maximum and minimum of the bootstrapped values. The dashed lines represent the 5% and 95 percentiles and the dotted line denotes the median.

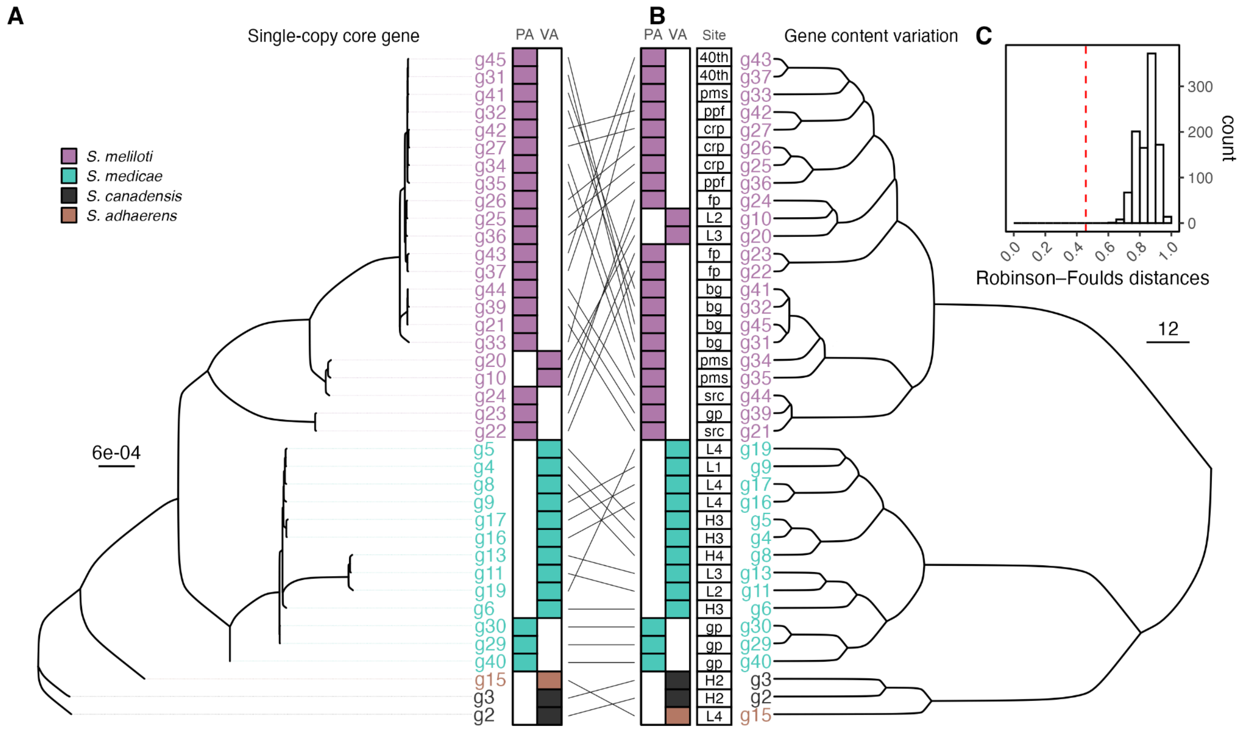


**Figure S4. Congruence of core and accessory gene phylogeny.** Panel (A) shows the phylogeny inferred from 987 single-copy core genes. The single-copy core-gene phylogeny represents the consensus tree of 1000 bootstrapped trees using the maximum likelihood approach. The branches with < 95% bootstrap support are collapsed. The internal nodes marked with gray shades represent a 1/100 scaling to branch length. (B) denotes the hierarchical clustering (ward.D2) of gene content variation inferred from the presence and absence of 26544 genes. The heatmap represents the isolates' origin in the Philadelphia (PA) or Virginia (VA) region. (C) The core and accessory phylogeny shown in panels A and B are more congruent than random expectation. The red line denotes observation and the histogram represents 1000 randomizations.

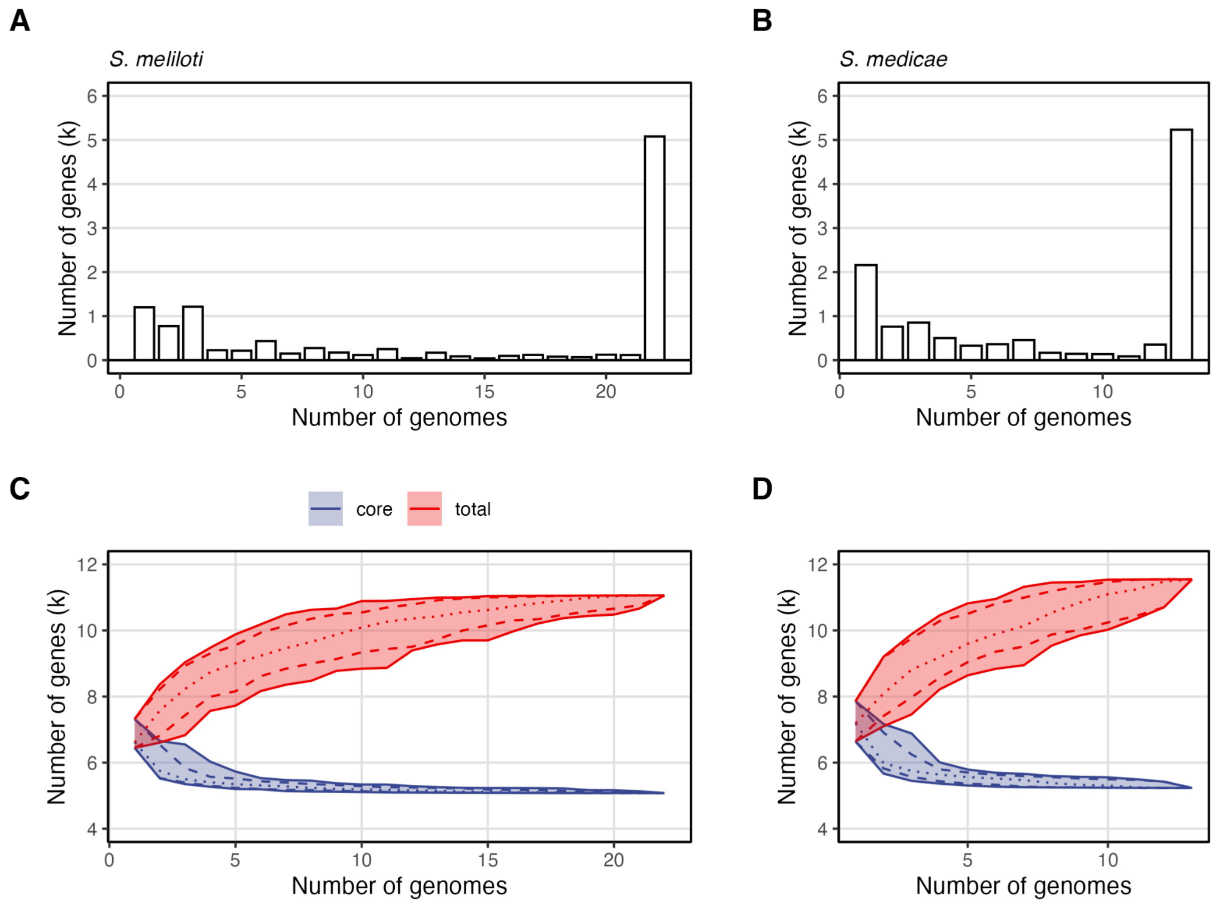


**Figure S5. Pangenome sampling within symbiotic species.** Panels A and C show the gene frequency spectrum and pangenome sampling in *S. meliloi*, respectively; Panels B and C show the corresponding figures in *S. medicae*. At each number of genomes, we performed 100 bootstraps and computed the number of core genes and the number of total genes. The solid lines represent the maximum and minimum of the bootstrapped values. The dashed lines represent the 5% and 95 percentiles and the dotted line denotes the median.

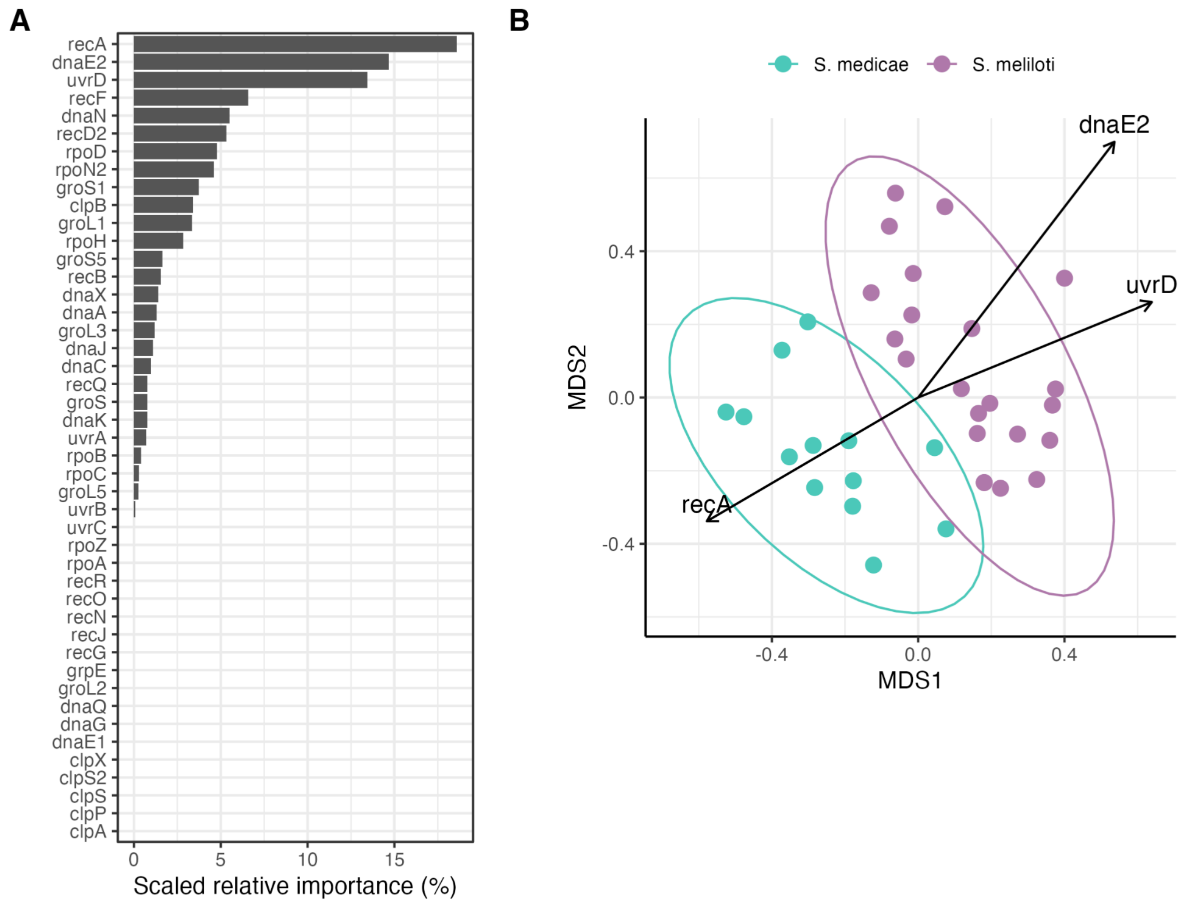


**Figure S6. Random forest and ordination analyses of heat-relevant accessory genes differentiating *S. meliloti* and *S. medicae*.** (A) Random forest variable importance scores (MeanDecreaseGini) were scaled to percentages to show the relative contribution of each gene to species classification. (B) Principal coordinates analysis (PCoA) based on Bray–Curtis dissimilarity of heat-relevant genes, with species shown by ellipses. Vectors from *envfit* indicate the direction and strength of key genes (recA, uvrD, dnaE2), showing that uvrD and dnaE2 are enriched toward *S. meliloti*, whereas recA is more associated with *S. medicae*.

### SUPPLEMENTARY TABLES

**Table S1. *Sinorhizobium* reference genomes.** Contigs in our genomes are assigned to one of four types based on their top BLAST hit: chromosome, pSymA, pSymB, and pAcce.

| | Accession | Species | Strain | pSymA | pSymB | pAcce |
|---|---|---|---|---|---|---|
| 1 | GCF_002197065.1 | *Sinorhizobium meliloti* | usda1106 | psymA | psymB | |
| 2 | GCF_000006965.1 | | em1021 | pSymA | pSymB | |
| 3 | GCF_013315775.1 | | em1022 | pA | pB | |
| 4 | GCF_002197445.1 | | usda1021 | psymA | psymB | accessoryA |
| 5 | GCF_000017145.1 | *Sinorhizobium medicae* | wsm419 | pSMED02 | pSMED01 | pSMED03 |
| 6 | GCF_021052565.1 | | wsm1115 | pWSM1115_2 | pWSM1115_1 | pWSM1115_3 |
| 7 | GCF_025200695.1 | | su277 | pSU277_2 | pSU277_1 | pSU277_3, pSU277_4 |
| 8 | GCF_000697965.2 | *Sinorhizobium adhaerens* | casidaa | | | |
| 9 | GCF_009883655.1 | | corn53 | | | |
| 10 | GCF_020035495.1 | | w2a | | | |
| 11 | GCF_020035515.1 | | w2b | | | |
| 12 | GCF_013488225.1 | *Sinorhizobium mexicanum* | ittgr7 | | | |
| 13 | GCF_017488845.2 | *Sinorhizobium canadensis* | t173 | | | |
| 14 | GCF_000018545.1 | *Sinorhizobium fredii* | ngr234 | | | |
| 15 | GCF_000219415.3 | | gr64 | | | |
| 16 | GCA_000283895.1 | | hh103 | | | |
| 17 | GCF_000705595.2 | *Sinorhizobium americanum* | ccgm7 | | | |
| 18 | GCF_002288525.1 | *Sinorhizobium sojae* | ccbau05684 | | | |
| 19 | GCF_008932245.1 | *Sinorhizobium alkalisoli* | yic4027 | | | |

Hijmans, Robert J. 2024. "Geosphere: Spherical Trigonometry."

Hijmans, Robert J., Marcia Barbosa, Aniruddha Ghosh, and Alex Mandel. 2023. "Geodata: Download Geographic Data." *R Package*.

Hufkens, Koen, David Basler, Tom Milliman, Eli K. Melaas, and Andrew D. Richardson. 2018. "An Integrated Phenology Modelling Framework in R: Modelling Vegetation Phenology with phenor." *Methods in Ecology and Evolution / British Ecological Society* 9:1–10.

Katoh, Kazutaka, Kazuharu Misawa, Kei-Ichi Kuma, and Takashi Miyata. 2002. "MAFFT: A Novel Method for Rapid Multiple Sequence Alignment Based on Fast Fourier Transform." *Nucleic Acids Research* 30 (14): 3059–66.

Kolmogorov, Mikhail, Jeffrey Yuan, Yu Lin, and Pavel A. Pevzner. 2019. "Assembly of Long, Error-Prone Reads Using Repeat Graphs." *Nature Biotechnology* 37 (5): 540–46.

Li, Heng. 2016. "Minimap and Miniasm: Fast Mapping and de Novo Assembly for Noisy Long Sequences." *Bioinformatics (Oxford, England)* 32 (14): 2103–10.

*Medaka: Sequence Correction Provided by ONT Research*. 2018. Github. https://github.com/nanoporetech/medaka.

Pebesma, Edzer. 2018. "Simple Features for R: Standardized Support for Spatial Vector Data." *The R Journal* 10 (1): 439.

Pebesma, Edzer, and Roger Bivand. 2023. *Spatial Data Science: With Applications in R*. New York: Chapman and Hall/CRC.

Reeve, Wayne, Patrick Chain, Graham O'Hara, Julie Ardley, Kemanthi Nandesena, Lambert Bräu, Ravi Tiwari, et al. 2010. "Complete Genome Sequence of the Medicago Microsymbiont Ensifer (Sinorhizobium) Medicae Strain WSM419." *Standards in Genomic Sciences* 2 (1): 77–86.

Seemann, T. n.d. "Barrnap 0.9 : Rapid Ribosomal RNA Prediction." https://github.com/tseemann/barrnap.

Seemann, Torsten. 2014. "Prokka: Rapid Prokaryotic Genome Annotation." *Bioinformatics (Oxford, England)* 30 (14): 2068–69.

Simão, Felipe A., Robert M. Waterhouse, Panagiotis Ioannidis, Evgenia V. Kriventseva, and Evgeny M. Zdobnov. 2015. "BUSCO: Assessing Genome Assembly and Annotation Completeness with Single-Copy Orthologs." *Bioinformatics (Oxford, England)* 31 (19): 3210–12.

Thornton, M. M., R. Shrestha, Y. Wei, P. E. Thornton, and S-C Kao. 2022. "Daymet: Daily Surface Weather Data on a 1-Km Grid for North America, Version 4 R1." ORNL Distributed Active Archive Center. https://doi.org/10.3334/ORNLDAAC/2129.

Tonkin-Hill, Gerry, Neil MacAlasdair, Christopher Ruis, Aaron Weimann, Gal Horesh, John A. Lees, Rebecca A. Gladstone, et al. 2020. "Producing Polished Prokaryotic Pangenomes with the Panaroo

Pipeline." *Genome Biology* 21 (1): 180.

Walker, Kyle. 2024. *Tigris: Download and Use Census TIGER/Line Shapefiles in R*. Github. https://github.com/walkerke/tigris.

Weidner, Stefan, Anke Becker, Ildefonso Bonilla, Sebastian Jaenicke, Javier Lloret, Isabel Margaret, Alfred Pühler, et al. 2012. "Genome Sequence of the Soybean Symbiont Sinorhizobium Fredii HH103." *Journal of Bacteriology* 194 (6): 1617–18.

Wickham, Hadley. 2016. *Ggplot2: Elegant Graphics for Data Analysis*. PDF. 2nd ed. Use R! Basel, Switzerland: Springer International Publishing. https://ggplot2.tidyverse.org.

Wick, Ryan R., and P. Menzel. 2017. "Filtlong." *Available Online: Github. com/rrwick/Filtlong (accessed on 15 August 2021)*. github.com/rrwick/Filtlong.

Williams, Laura E., David A. Baltrus, Sean D. O'Donnell, Tara J. Skelly, and Mark O. Martin. 2017. "Complete Genome Sequence of the Predatory Bacterium Ensifer Adhaerens Casida A." *Genome Announcements* 5 (47). https://doi.org/10.1128/genomeA.01344-17.

Wood, Simon N. 2011. "Fast Stable Restricted Maximum Likelihood and Marginal Likelihood Estimation of Semiparametric Generalized Linear Models: Estimation of Semiparametric Generalized Linear Models." *Journal of the Royal Statistical Society. Series B, Statistical Methodology* 73 (1): 3–36.
